# Effects of food restriction on body mass, energy metabolism and thermogenesis in a tree shrew (*Tupaia belangeri*)

**DOI:** 10.1101/478487

**Authors:** Xue-na Gong, Hao Zhang, Di Zhang, Wan-long Zhu

## Abstract

This study investigates the energy strategies of a small mammal in response to food shortages as a function of food restriction (FR), metabolic rate and ambient temperature. We subjected tree shrews (*Tupaia belangeri*) to FR and measured body mass, survival rate, resting metabolic rate (RMR), nonshivering thermogenesis (NST) and cytochrome *c* oxidase (COX) activity of brown adipose tissue (BAT). Cold-exposed animals restricted to 80% of *ad libitum* food intake had significantly increased RMR and NST and decreased body mass and survival rates compared with those kept at room temperature on the same FR level. Animals classified has having a high RMR consumed 30.69% more food than those classified as having a low RMR, but showed no differences in body mass or survival when restricted to 80% of *ad libitum* food intake. These results indicate that tree shrews, known for their relatively high metabolic rates, are sensitive to periods of FR, which supports the metabolic switch hypothesis. Our findings are also consistent with the prediction that small mammals with food hoarding behaviors, like tree shrews, may have a lower tolerance for food shortages than non-hoarding species.

## Introduction

Adaptive regulation of energy metabolism plays an important role in small mammals for coping with changes in the natural environment. In nature, many animals are faced with seasonal variations in food resources (Veloso and Bozinovic 1993). Non-hibernating small mammals generally respond to a food shortage in one of two ways: either by reducing their energy metabolism levels (e.g. MF1 mice (Hambly and Speakman 2005), white-footed mouse (*Peromyscus maniculatus*) (Gutman et al. 2007), chevrier’s field mouse (*Apodemus chevrieri*) (Zhu et al. 2013) and yunnan red-backed vole (*Eothenomys miletus*) (Zhu et al. 2014); or by maintaining or increasing their energy expenditure levels (e.g. KM mice, rats and guinea pigs (*Cavia porcellus*) (Williams et al. 2002; Zhao et al. 2009*a*, 2009*b*)). According to the metabolic switch hypothesis, an ability to adjust metabolic rate plays a key role in animals adapted to periods of food shortage, enabling them to “switch down” their resting metabolic rate to survive these shortages (Merkt and Taylor 1994). The adaptive strategies adopted by animals in response to food shortages are also affected by other environmental conditions and their own behavior. In a 50% food restriction (FR) experiment, the golden thorn rat hair (*Acomys russatus*), a non-hoarding species, survived more than 6 weeks, while the foxtail gerbil (*Gerbillus dasyurus*), a hoarding species, survived only 2 weeks. These data suggest that hoarding species may have a lower tolerance to food shortages than non-hoarding species (Gutman et al. 2006). Whether this is generally true, or whether the survival of hoarding species during food shortages also depends on their capacity for metabolic rate adjustment, remains unclear.

The tree shrew *Tupaia belangeri* (Mammalia: Scandentia: Tupaiidae) is a small mammal native to Southeast Asia. It is widely distributed throughout Southern China, India and Southeast Asia in farmland and shrub habitat. Tree shrews are omnivorous, eating mainly insects, melons and fruits, and grains, and are a food hoarding species. Their food supply is affected by seasonal changes, and their diets reflect these changes (Wang et al. 1991). It has been reported that tree shrews show seasonal thermogenesis (Zhu et al. 2012) and have a higher resting metabolic rate (RMR) in their thermoneutral zone (TNZ) than other small mammals (Wang et al. 1994; Xiao et al. 2010). This may be associated with food resource and quality, as fasting and FR has been demonstrated to significantly decrease metabolic thermogenesis (Gao et al. 2014, 2016). Food supply may thus have influenced the evolution of metabolic plasticity in tree shrews. The aim of this study was to test the metabolic switch hypothesis in tree shrews by investigating their energy strategies in response to food shortages as a function of metabolic rate and ambient temperature and measured changes in body mass, metabolic thermogenesis and cytochrome *c* oxidase (COX) activity of brown adipose tissue (BAT).

## Materials and Methods

### Experimental animals

Tree shrews, *T. belangeri*, were wild-captured from farmland and shrub near Luquan County (25°26′–25°62′ N, 102°13′–102°57′ E, altitude 1650–1700 m) in Yunnan Province, China. The area is located in the northern part of the Yunnan Plateau and has a subtropical plateau climate. The geological landform is complex with large variations in surface relief. The temperature shows marked changes with altitude, and while the annual temperature changes are small, there are large daily temperature fluctuations. Tree shrews were transported to the Animal Feeding Room of Yunnan Normal University, and individually housed in wire cages (40×30×30 cm). Animals were kept in a room maintained at 25 ± 1 °C with a natural photoperiod for 4 weeks prior to the experiment, and provided *ad libitum* food and water. The food contained the following ingredients (by weight): 5% milk, 5% sugar, and 90% cornmeal. All procedures were licensed under the Animal Care and Use Committee of the School of Life Science, Yunnan Normal University (approval ref. 13-0901-011).

#### Experiment 1: Effects of FR and low temperature on body mass, energy metabolism and thermogenesis

We randomly divided 20 adult tree shrews of similar body mass into two groups (N = 10 per group): Warm (room temperature, 25 □ ± 1 □) and Cold (low temperature, 5 □ ± 1 □). Animals were acclimated to their respective temperatures with *ad libitum* food intake for 4 weeks (d28–d0), and then restricted to 80% of initial *ad libitum* food intake for 5 days (d1–d5). Animal survival rate and body mass were measured daily, and the RMR, NST and COX activity of BAT were measured on the 6th day.

##### Metabolic trials

Metabolic rates were measured using an AD ML870 open respirometer at 30 ± 0.5 °C within the TNZ as described previously (the TNZ of *T. belangeri* is 30–35 °C, Wang et al. 1994). The volume of the metabolic chamber was 760 ml and the temperature in the chamber was maintained within 0.5 °C by a SPX-300 temperature-controlled cabinet. The air flow was 200 ml/min, and a ML206 gas analyzer was used for gas analysis. Tree shrews were fasted for 3 h before being transferred into the metabolic chamber. After 1 h of adaptation to the chamber, metabolic measurements were conducted for another 1 h, during which oxygen consumption was read at 10 s intervals. RMR was calculated using two continuous, stable, minimum recordings (Wang et al. 1994) using the method of Hill (1972). Nonshivering thermogenesis (NST) was induced by a subcutaneous injection of norepinephrine (Shanghai Harvest Pharmaceutical) and measured at 30 °C. The dose of norepinephrine administered was approximately 0.8–1.0 mg/kg body mass, used based on dose-dependent response curves generated before the experiment (Zhu et al. 2008*a*). NST was calculated using the two highest consecutive recordings of oxygen consumption (Wang et al. 1995).

##### BAT mass and COX activity

Interscapular BAT was removed and weighed immediately after the experiment. Mitochondrial protein (MP) was prepared as described in a previous report (Wiesinger et al. 1989). MP content was determined by the Folin phenol method (Lowry et al. 1951), with bovine serum albumin as the standard. The COX activity of BAT was measured polarographically with oxygen electrode units (Hansatech Instruments, England; Sundin et al. 1987).

#### Experiment 2: Effects of FR and metabolism levels on body mass, energy metabolism and thermogenesis

Thirty adult tree shrews with similar body mass were breeding 4 weeks before experiment. We measured the RMR of each animal, and assigned them to one of two groups accordingly (N = 10 per group): hRMR (higher RMR) and lRMR (lower RMR). There was a significant difference in RMR between these two groups (*t* = 5.428, *P* < 0.01). *Ad libitum* food intake was measured for each group, and food was then restricted to 80% of this intake for 14 days. Animal survival rate and body mass were measured daily. Interscapular BAT and liver were removed immediately after the experiment, and the COX activity of BAT and liver were measured using the methods outlined in experiment 1.

Animal wet carcass mass was recorded after the visceral organs and digestive tract were removed. The carcass was then dried in an oven at 60 °C to constant mass and reweighed to get the dry mass. Body fat was extracted from the dried carcass by ether extraction in a Soxtec2043 apparatus. Body fat content (%) was calculated as body fat mass/dry carcass mass × 100% (Zhao and Wang 2006).

##### Statistical analysis

Data were analyzed using the SPSS 16.0 software package. Differences in RMR, NST, MP content and COX activity of BAT or liver, carcass mass and body fat content were examined using one-way ANOVA or ANCOVA with body mass or carcass mass as a covariate, followed by least-significant difference (LSD) post-hoc tests. Differences between groups on a single experimental day were examined using independent t-tests. Results are presented as means ± SE and *P* < 0.05 was considered to be statistically significant.

## Result

### Experiment 1

#### Survival rate, body mass and food intake

After 5 days of FR-80%, the survival rates of Warm and Cold were 100% and 60%, respectively (fig. 1*A*). Temperature also had significant effects on the rate of change in body mass. By day 5, body mass of FR-80% was reduced by 4.64% in Warm, and 17.41% in Cold (*F*_5,25_ = 17.541, *P* < 0.01, fig. 1*B*). Food intake was also affected by temperature, with Cold FR-80% consuming more food than Warm FR-80% (d0, *t* = 5.634, *P* < 0.01; d5, *t* = 5.357, *P* < 0.01, fig. 1*C*).

**Fig. 1.**
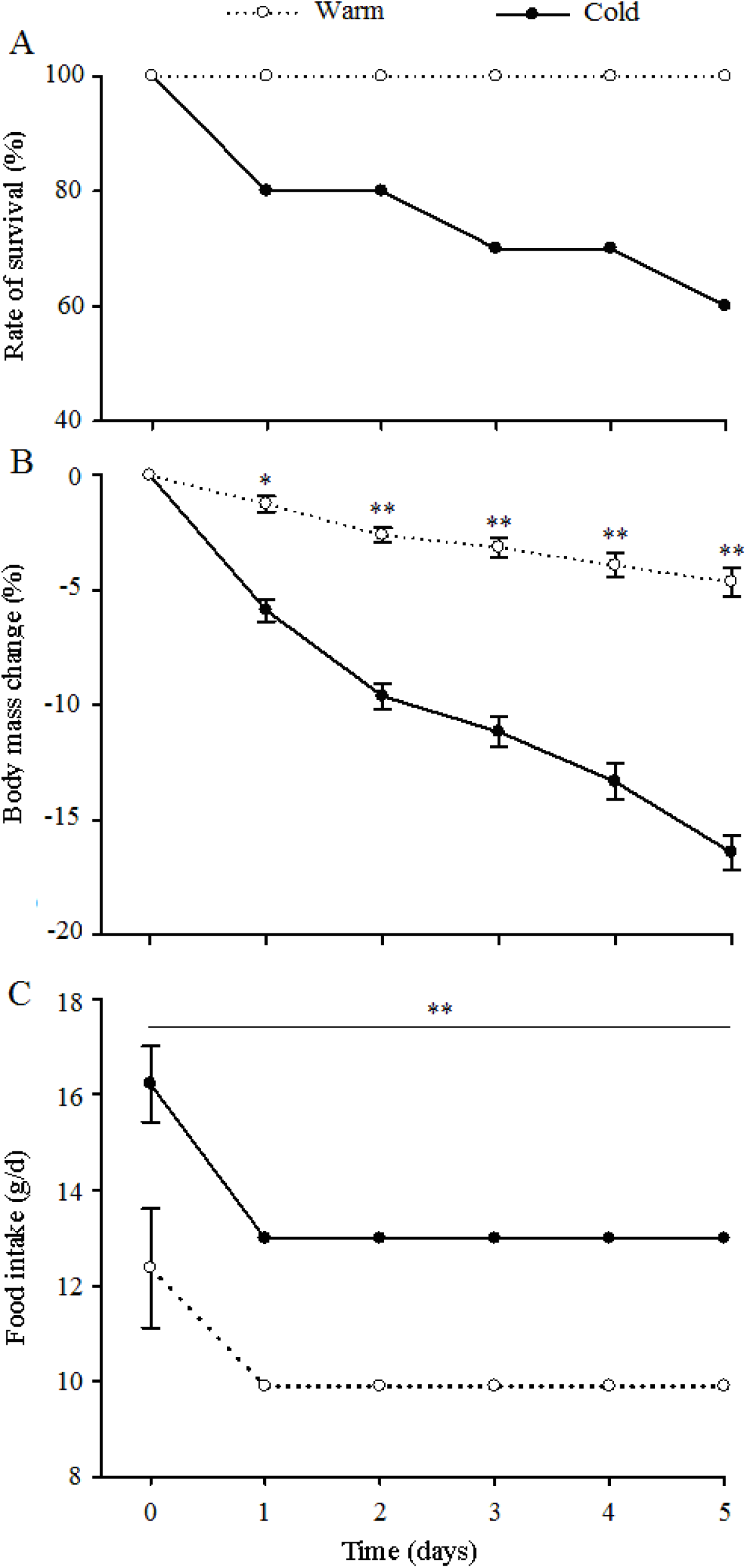
Effect of ambient temperature on the rate of survival (A), body mass change (B) and food intake (C) in tree shrews restricted to 80% of *ad libitum* food intake. Warm, 25°C, Cold, 5°C, *, *P*< 0.05, **, *P* < 0.01

#### RMR, NST and BAT COX activity

RMR and NST were increased after cold acclimation for 4 weeks, and were significantly higher at 4 weeks than those of the room temperature group (RMR, *t* = 6.415, *P* < 0. 01; NST, *t* = 7.236, *P* < 0.01, fig. 2). After 5 days of FR-80%, RMR and NST of Cold remained significantly higher than those of Warm (RMR, 20.92%, *t* = 5.861, *P* < 0.01; NST, 24.73%, *t* = 6.135, *P* < 0.01, fig. 2). BAT mass, MP content and COX activity of BAT of Cold FR-80% were also significantly higher in than Warm FR-80% (BAT, *t* = 4.137, *P* < 0.01, fig. 3*A*; MP content, *t* = 3.542, *P* < 0.01, fig. 3*B*; COX activity, *t* = 3.376, *P* < 0.05, fig. 3*C*).

**Fig. 2.**
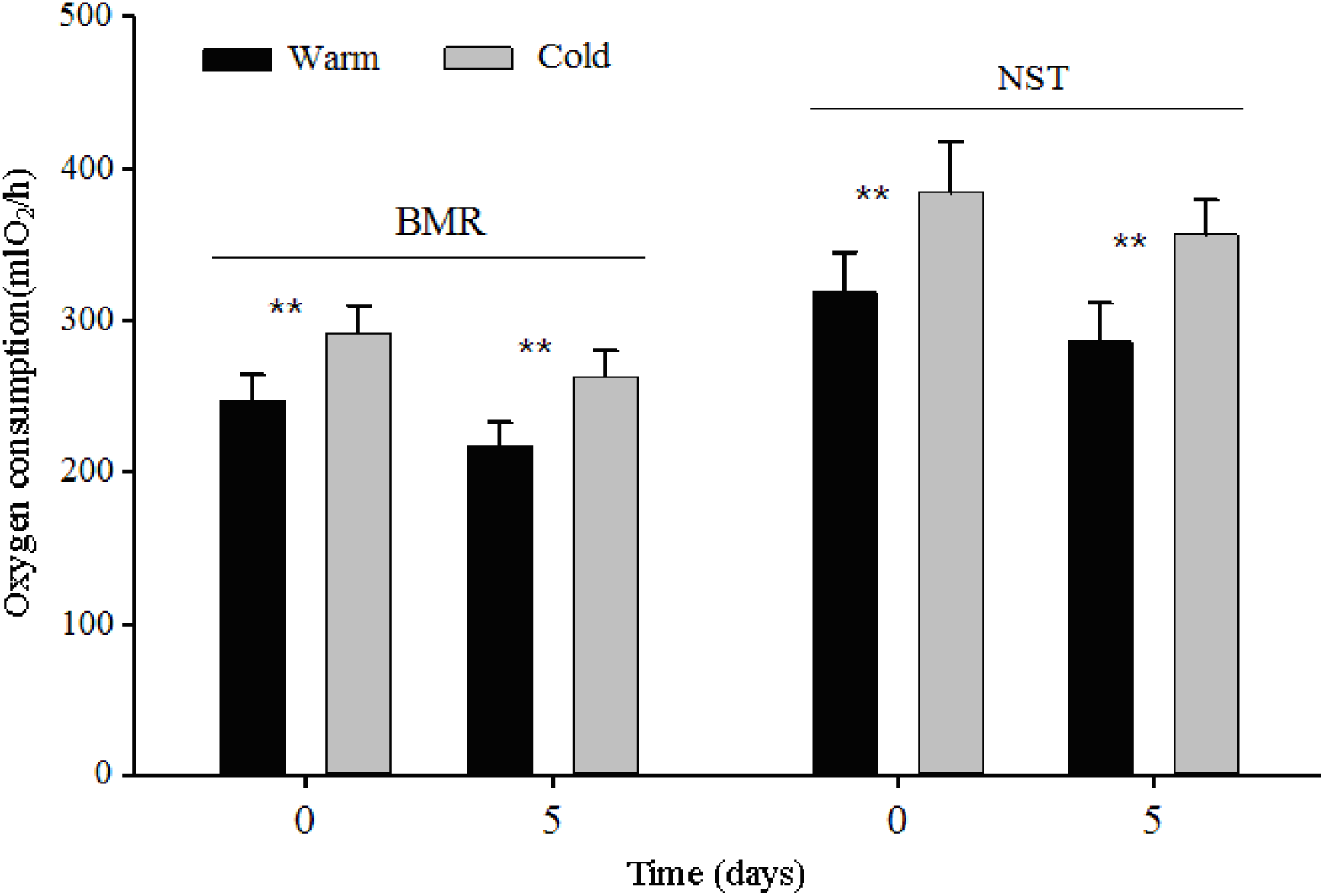
BMR and NST in tree shrews that were exposed to cold for 4 weeks (d0), and then were restricted to 80% of *ad libitum* food intake for 5 days (d5). Warm, 25°C, Cold, 5°C, **, *P* < 0.01

**Fig. 3.**
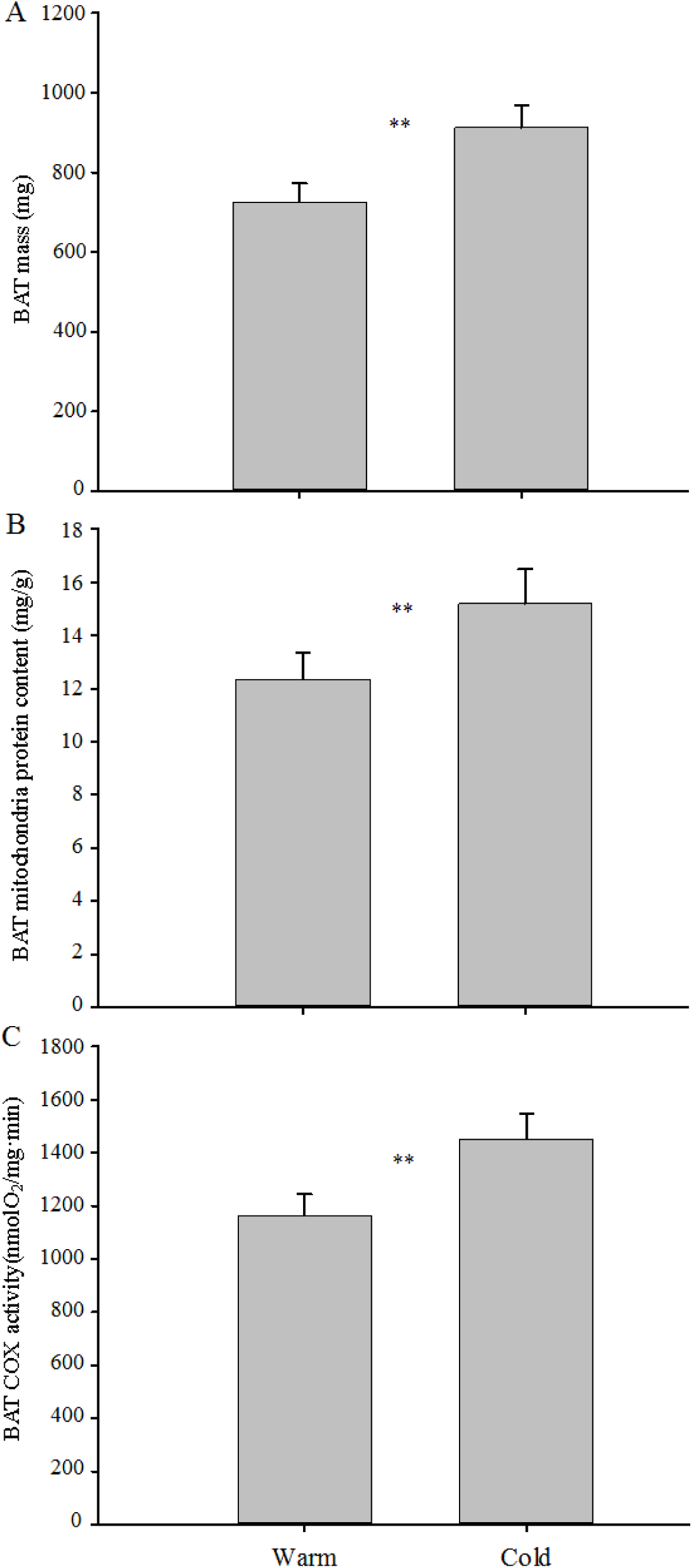
BAT mass (A), mitochondria protein content (B) and COX activity (C) of BAT in tree shrews exposed to cold for 4 weeks and then were restricted to 80% of ad libitum food intake for 5 days. Warm, 25°C, Cold, 5°C, **, *P* < 0.01

### Experiment 2

#### Survival rate, body mass and food intake

Prior to FR, there was a significant difference in the food intake of hRMR and lRMR, with hRMR consuming 30.69% more food than the lRMR (*t* = 5.443, *P* < 0.01, fig. 4*A*). On FR day 14, the survival rate of hRMR was 70% and lRMR was 80% (fig. 4*B*). During the period of FR, the body mass of both groups significantly decreased, but the difference between the groups was not significant (*t* = 1.113, *P* > 0.05, fig. 4*C*).

**Fig. 4.**
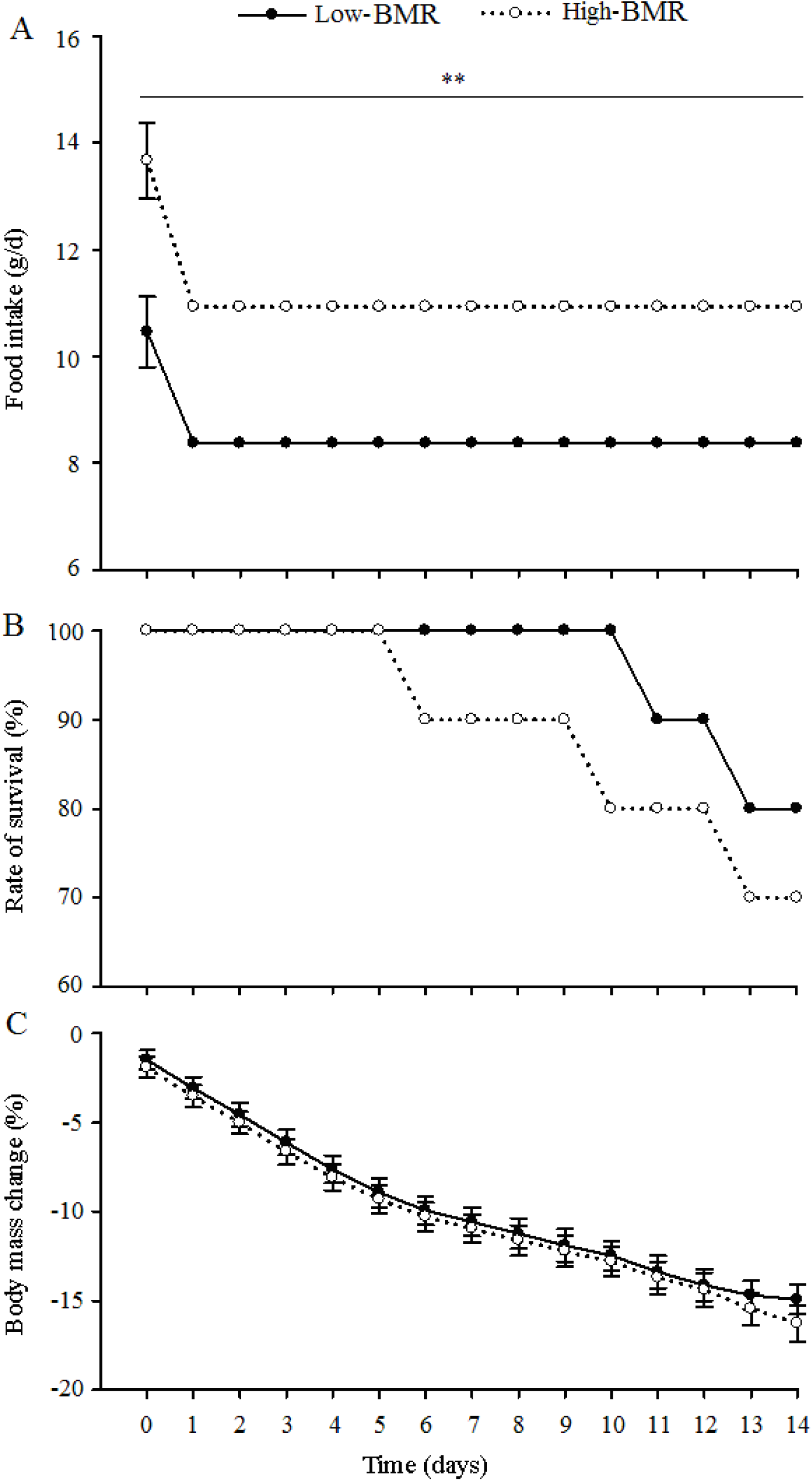
Food intake (A), rate of survival (B) and body mass change (C) in tree shrews with high- or low-BMR restricted to 80% of *ad libitum* food intake. **, *P* < 0.01

#### BMR, NST, carcass mass, body fat content, COX activity of BAT and liver

RMR was significantly higher in hRMR than lRMR (*t* = 5.428, *P* < 0.01). During the 14-day FR period, RMR significantly decreased in hRMR but showed no significant change in lRMR. On FR day 14, RMR remained significantly higher in hRMR than lRMR (Table 1). There were no significant differences between groups in NST, wet or dry carcass mass, body fat content, or COX activity of BAT or liver (Table 1).

**Table 1.** Effect of food restriction on the rate of metabolism and activity of cytochrome c oxidase (COX) of brown adipose tissue (BAT) and liver in tree shrews with high-or low-BMR

|  | Low BMR | High BMR | *P* |
| --- | --- | --- | --- |
| Body mass (g) | 108.41±3.15 | 107.53±3.21 | ns |
| Wet carcass mass (g) | 90.36±2.87 | 88.69±3.11 | ns |
| Dry carcass mass (g) | 41.41±2.39 | 38.53±3.54 | ns |
| Body fat content (%) | 25.72±1.94 | 25.54±1.86 | ns |
| BMR (mlO<sub>2</sub>/h) | 206.44±13.73 | 241.62±15.33 | $P < 0.05$ |
| NST (mlO<sub>2</sub>/h) | 317.73±26.35 | 328.28±24.19 | ns |
| BAT |  |  |  |
| Mass (g) | 0.55±0.04 | 0.59±0.03 | ns |
| Mt protein content (mg/g tissue) | 10.12±0.67 | 10.48±0.76 | ns |
| Mt protein content (mg/total tissue) | 5.57±0.44 | 6.18±0.48 | ns |
| COX activity (nmolO<sub>2</sub>/min·mg Mt protein) | 941.23±52.36 | 1006.54±64.57 | ns |
| COX activity (nmolO<sub>2</sub>/min·total tissue) | 1139.64±62.34 | 1218.49±76.51 | ns |
| Liver |  |  |  |
| Mass (g) | 4.88±0.21 | 4.83±0.18 | ns |
| Mt protein content (mg/g) | 30.24±2.58 | 33.37±2.16 | ns |
| Mt protein content (mg/total tissue) | 148.57±4.11 | 163.01±3.76 | ns |
| COX activity (nmolO<sub>2</sub>/min·mg Mt protein) | 82.63±2.67 | 86.46±3.54 | ns |
| COX activity (nmolO<sub>2</sub>/min-total tissue) | 3124.75±124.56 | 3428.66±137.25 | ns |

Data are means±SE. ns: non-significant ( $P > 0.05$ )

## Discussion

Temperature is one of the most important environmental factors affecting metabolic rate (Wang et al. 1999; Wang et al. 2015), Small mammals acclimated to low temperature environments have higher metabolic rates (Klingenspor 2003; Tang et al. 2009; Chi and Wang 2011). In this study, cold acclimation significantly increased the metabolic rate of tree shrews, similar to Zhang et al. (2012*a*, *b*). In response to the increase in energy expenditure required at low temperature, tree shrews increased their food intake by 24.01%. However, after 5 days of FR-80% at low temperature, body mass decreased by 17.41% and survival rate dropped to 60%. These findings indicate that the increase in metabolism in response to low temperatures reduced their tolerance to FR. This supports the notion that the relatively high metabolic rates of tree shrews may underpin their low tolerance to FR.

Studies of small mammals have shown both inter- and intraspecific differences in metabolic rates. Here, we found significant differences in RMR among individual tree shrews. Animals from the high RMR group consumed 30.69% more food than those from the low RMR group, and significantly reduced their metabolic rate in response to FR. Similarly, Sundevall’s jird (*Meriones crassus*) and the golden spiny mouse (*Acomys russatus*) have been shown to reduce their metabolic rate in response to FR, also supporting the metabolic switch hypothesis (Merkt and Taylor 1994; Gutman et al. 2006, 2007). However, after 2 weeks of FR-80%, the survival rate of the high RMR group from our study dropped to 70%, suggesting that they did not establish new energy balance through reduced metabolism in response to FR. The metabolic rates of the low RMR group were relatively low, and their food intake was significantly lower than that of the high RMR group. The RMR of the low RMR group did not significantly decrease during FR, suggesting that energy intake was insufficient to compensate for energy expenditure and animals were in a state of negative energy balance. These findings further indicate that the relatively high metabolic rates of tree shrews underpin their low tolerance of FR relative to their food intake and body size.

*T. belangeri* have the northern-most distribution of all Southeast Asian tree shrews (Wang et al. 1991), found mainly on the Yunnan-Guizhou Plateau, where the animals for this study were captured. Their habitat has a subtropical plateau monsoon climate, with more low temperature stress than a tropical climate (Wang et al. 1994). *T. belangeri* have a higher metabolic rates than other tree shrews, such as the Philippine tree shrew (*Urogale everetti*), common tree shrew (*Tupaia glis*) and pen-tailed tree shrew (*ptilocercus lowii*) (Wang et al. 1994). High metabolic rates are advantageous in adapting to cold climate conditions in winter (Zhu et al. 2012) and may also relate to their solitary behavior, which typically demands more energy for temperature regulation than do social behaviors (McNab 1988; Song and Wang 2003). However, the degree of increase in RMR in response to low temperatures is lower in tree shrews than the yunnan red-backed vole (*Eothenomys miletus*) by region (Zhu et al. 2008b). The effect of low temperature on RMR in small mammals is related to the geographical distribution of their species. Thus, RMR plays an important role in the tree shrews’ adaptive response to low temperature stress. It also suggests that low temperature may be one of the factors preventing tree shrews from extending their range further north.

In summary, under natural conditions, tree shrews faced with seasonal changes in food resources have a lower tolerance to food shortages than other small mammals. This is consistent with the prediction that small mammals that hoard food are less able to cope with food shortages than non-hoarding species. We suggest that the main reason for the tree shrews’ low tolerance to food shortages is their high metabolic rate relative to food intake and body size. Tree shrews appear to maintain energy balance through adjustments in thermogenesis, and this metabolic plasticity likely evolved as an adaptive strategy to food shortages.

## Acknowledgments

This research was financially supported by National Science Foundation of China (No. 31660121), The Yunnan Provincial Key Program for Applied Basic Research (No. 2016FA045).

